# Hypofucosylation promotes pertussis toxin binding to cell surface glycococonjugates and pertussis toxin-induced intracellular ERK signaling

**DOI:** 10.64898/2025.12.03.692184

**Authors:** Rohit Sai Reddy Konada, Nicole Nischan, Aurora Silva, Jennifer J. Kohler

## Abstract

Pertussis (whooping cough) is caused by the bacterium *Bordetella pertussis*. Among the virulence factors produced by *B. pertussis,* pertussis toxin (PT) is responsible for key disease symptoms, including impacts on the immune system. PT is an AB_5_ toxin, consisting of a single catalytic A subunit and five B subunits. The B pentamer recognizes cell surface glycans and facilitates intracellular delivery of the catalytic A subunit. PT also impacts host cell signaling via mechanisms that do not depend on the catalytic activity of the A subunit. In particular, PT promotes signaling through the T-cell receptor (TCR), leading to activation of extracellular signal-regulated kinase (ERK) cascades. PT prefers to bind sialylated and N-linked glycans, but other aspects of PT’s glycan binding specificity remain underexplored. Here we report that the absence of fucose on mammalian cell surfaces leads to increased binding by PT. Using pharmacological inhibitors in a human bronchial epithelial cell line, we observe that sialylation and N-linked glycosylation promote PT binding while fucosylation interferes with PT binding. Similarly, CHO and Colo205 cells deficient in fucosylation exhibited enhanced PT binding as compared to the corresponding wild-type cell lines. Genetic knockout of FUT3/FUT5/FUT6 or of FUT8 led to increased PT binding, suggesting that specific fucosylated epitopes mediate protection from PT. The functional impact of altered PT binding was examined in Jurkat T cells, where removal of cell surface non-core fucose led to increased PT-dependent ERK phosphorylation. In sum, our study identifies a role for fucosylation in protecting mammalian cells from PT.

## INTRODUCTION

Pertussis (whooping cough) is a severe respiratory disease caused by a human-specific pathogenic bacterium *Bordetella pertussis*. Pertussis typically affects infants more severely compared to other age groups.(1) Symptoms in infected infants include paroxysmal cough with whooping sound, vomiting, pulmonary complications, and rib fractures.(2,3) *B. pertussis*, being a strict human pathogen without known reservoirs, transmits through respiratory droplets.(4) Genetic analysis of the pathogen combined with historical accounts reveals that pertussis disease has existed for centuries.(5) With consistent outbreaks, pertussis remains a major endemic disease in both developed and developing countries.(6) Introduction of a vaccine in 1940s drastically reduced the number of cases compared to the pre-vaccine era, but sporadic outbreaks remain common across the globe.(7) According to the Centers for Disease Control and Prevention, cases are on the rise in United States due to multiple factors including waning immunity.(8)

Pertussis results from a combined interplay of several virulence factors produced by *B. pertussis*, one of which is pertussis toxin (PT).(9) PT-deficient strains are extremely rare.(10,11) Conversely, increases in pertussis cases have been linked to emergence of strains with increased PT production.(12) Antibodies against PT are protective against severe disease and PT is a key component of acellular pertussis vaccines. Indeed, a mono-component acellular vaccine containing only detoxified PT has effectively controlled pertussis in Denmark.(13) The mechanisms by which PT contributes to disease are incompletely understood. Experiments in mice demonstrate that PT plays a role in establishing respiratory infection.(14) PT inhibits early chemokine production, causing impaired neutrophil recruitment.(15,16) The target cell type remains unclear: PT can inhibit chemokine production by macrophages but direct effects on epithelial cells have also been suggested.(15,17) PT is also implicated in leukocytosis,(18,19) a hallmark of severe disease that is proposed to occur through PT intoxication of lymphocytes, leading to reduced lymphocyte extravasation.(20) Finally, experiments performed in rat and baboon animal models point to a possible role for PT in the paroxysmal cough.(21,22) Taken together, existing data indicate that PT has multiple physiological effects stemming from action on distinct cell types, which may include lymphocytes, macrophages, respiratory epithelial cells, and nociceptor neurons.

PT is an AB_5_ toxin comprising a single catalytic A subunit (S1) and five B subunits (one S2, one S3, two S4, and one S5) that mediate binding to the surface of target cells. The holotoxin enters cells through endosomal uptake and is retrograde transported through the Golgi to the ER.(23,24) The S1 subunit enters the cytoplasm and ADP-ribosylates the α-subunit of heterotrimeric G proteins, resulting in dysregulation of intracellular signal transduction pathways.(25) However, some effects of PT, such as T cell mitogenicity, are independent of ADP-ribosylation activity.(26,27) PT binding to host cells depends on the S2 and S3 subunits, which have ∼70% sequence identity.(28,29) PT was co-crystallized with a sialylated N-linked glycopeptide from transferrin, revealing shallow binding pockets on the S2 and S3 subunits that can recognize terminal N-acetylneuraminic acid (Neu5Ac; form of sialic acid) residues.(30) Additionally, recent cryo-EM analysis of PT complexed with neutralizing antibodies yielded evidence for additional glycan binding sites on S2 and S3.(31) Sialic acid is commonly described to be the receptor for PT, but glycan array analysis and quantitative carbohydrate binding assays indicate that PT can also recognize non-sialylated glycans.(32–36) Additional insight into the glycan features important for PT affinity has been gleaned from in vitro binding assays.(35,36) However, binding affinity is not the only characteristic that defines physiologically relevant receptors: additional factors including glycoconjugate abundance and glycoconjugate clustering impact which cell surface ligands are most important for recognition by glycan-binding proteins.(37)

Here we examine glycosylation features that regulate PT binding to different cell types including two – respiratory epithelial cells and lymphocytes – that are possible physiological targets of PT. We confirm that PT preferentially binds sialylated and N-linked glycoconjugates. Additionally, we report that specific forms of fucosylation lead to reductions in PT binding. Further, we find that non-core fucosylation protects T-cell lymphocytes against PT-induced extracellular signal-regulated kinase (ERK) signaling. Together, these results provide insight into possible mechanisms by which variations in host glycosylation could modulate the severity of PT-induced impacts on the immune system.

## RESULTS

### Inhibition of fucosylation in HBEC-3KT cells results in increased PT binding

To model human airway epithelium, we used HBEC-3KT cells, an hTERT-immortalized bronchial epithelial cell line.(38) We modulated glycan biosynthesis in HBEC-3KT cells using pharmacological inhibitors of glycosyltransferases and glycosidases (Fig. 1A). We established appropriate inhibitor concentrations and incubation times by measuring the impact on cell surface binding of lectins and antibodies that recognize specific glycan structure (Supplementary Fig. 1). After pharmacological treatments, binding of PT to HBEC-3KT cell surfaces was analyzed by flow cytometry (Fig. 1B) and PT lectin blot (Fig. 1C). Culturing cells with kifunensine, an inhibitor of N-glycan maturation, resulted in significantly reduced PT binding in both assays (Fig. 1B and 1C), consistent with prior studies demonstrating that PT can bind N-linked glycans.(39,40) Culturing HBEC-3KT cells with sialylation inhibitor peracetylated 3-fluoro-β-D-N-acetylneuraminic acid (3F_ax_Neu5Ac)(41) also resulted in greatly reduced the PT binding in both assays (Fig. 1B and 1C), consistent with the well-documented preference of PT to bind sialylated glycoconjugates.(40,42) Next, we assessed the role of fucosylation in PT binding. HBEC-3KT cells cultured with peracetylated 2-deoxy-2-fluoro-L-fucose (2F-Fuc)(41,43) displayed increased PT binding as assessed by both flow cytometry and PT lectin blot. These results demonstrate that PT preferentially binds sialylated N-glycans in HBEC-3KT cells and suggest that PT prefers to bind non-fucosylated glycans.

**Figure 1.**
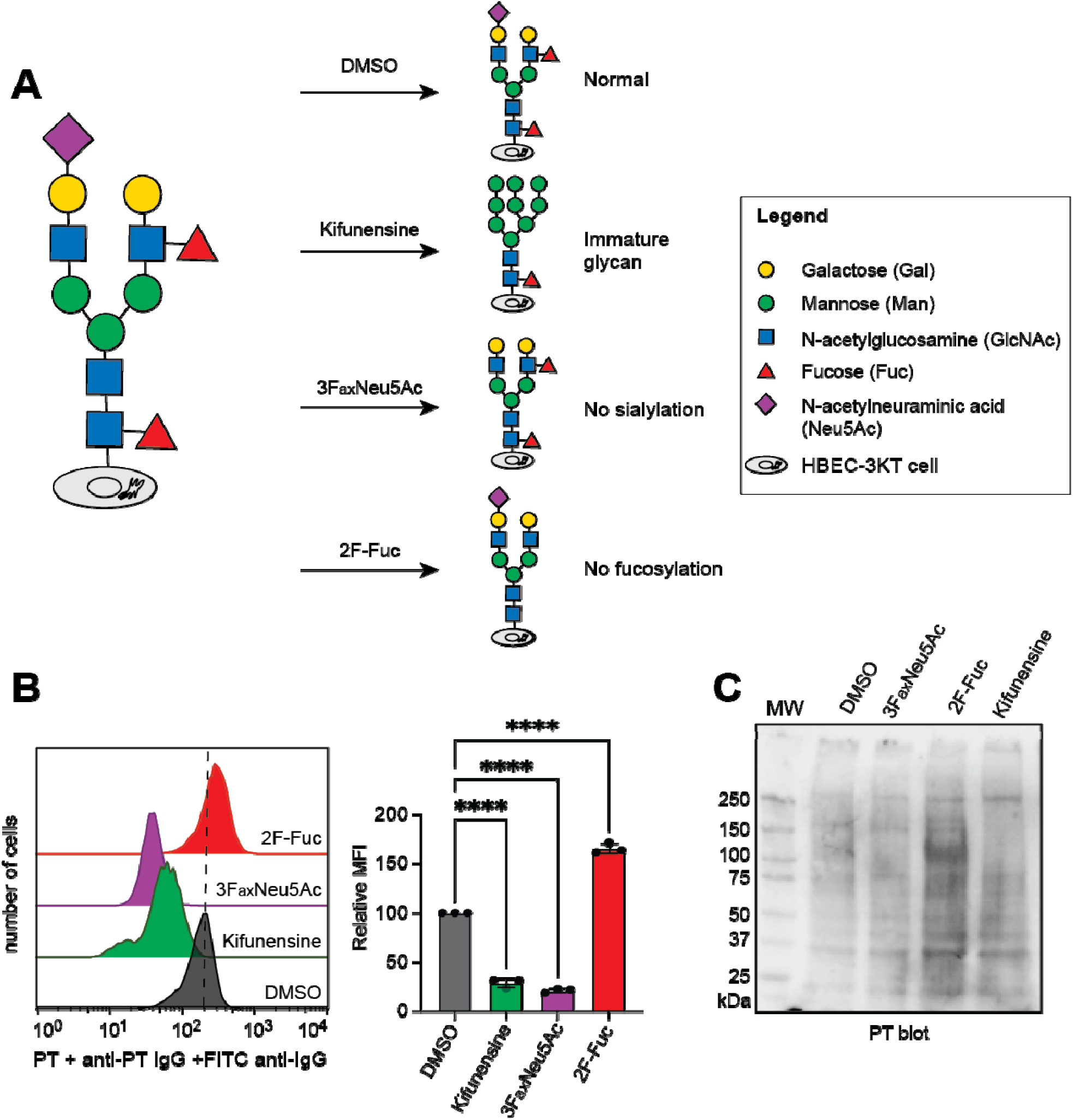
Inhibition of fucosylation in HBEC-3KT cells results in increased PT binding. **(A**) Schematic of glycan synthesis inhibition in HBEC-3KT cells using small molecule inhibitors of glycosylation. HBEC-3KT cells were cultured in the presence of the inhibitors for 72 h. **(B)** Flow cytometr analysis of PT binding to cell surfaces of the HBEC-3KT cells cultured with DMSO, kifunensine, 3F_ax_Neu5Ac, or 2F-Fuc. Bar graph shows the quantification of geometric mean fluorescence from 3 independent trials, normalized to the geometric mean fluorescence of DMSO-treated cells. Statistical analyses were performed by one-way ANOVA (Error bar indicates mean ± SD, **** indicates p value < 0.0001). **(C)** PT lectin blot of lysates of DMSO-, kifunensine-, 3F_ax_Neu5Ac-, or 2F-Fuc-treated HBEC-3KT cells. The blot shown is representative of 3 biological replicates.

### CHO cells devoid of fucose exhibit more PT binding than wild-type CHO cells

Chinese hamster ovary (CHO) cells exhibit a characteristic clustered growth pattern in response to incubation with PT.(44) This response is commonly used for the quantification of residual PT activity in acellular pertussis vaccines.(45) As CHO cells represent an important tool for evaluating PT activity, we therefore assessed which glycoconjugate features are required for PT binding to CHO cells. We obtained the CHO mutant cell line Lec2, which lacks sialylated glycoconjugates,(46–48) and Lec13 mutant, which lacks fucosylated glycoconjugates,(48–50) as well as the W5 parental CHO cell line (Fig. 2A). As expected, PT exhibited less binding to non-sialylated Lec2 CHO cell surfaces as compared to W5 cell surfaces (Fig. 2B). Similarly, while PT bound to glycoconjugates found in lysates from W5 cells, no binding was observed to Lec2 cell lysates (Fig. 2C). In contrast, PT exhibited more binding to non-fucosylated Lec13 CHO cell surfaces as compared to W5 cells (Fig. 2B). Additionally, lectin blot analysis showed more PT binding to Lec13 lysates as compared to W5 cell lysates (Fig. 2C). These results extend the observations made with HBEC-3KT cells, indicating that PT preferentially binds to glycoconjugates from cells that lack fucosylation.

**Figure 2.**
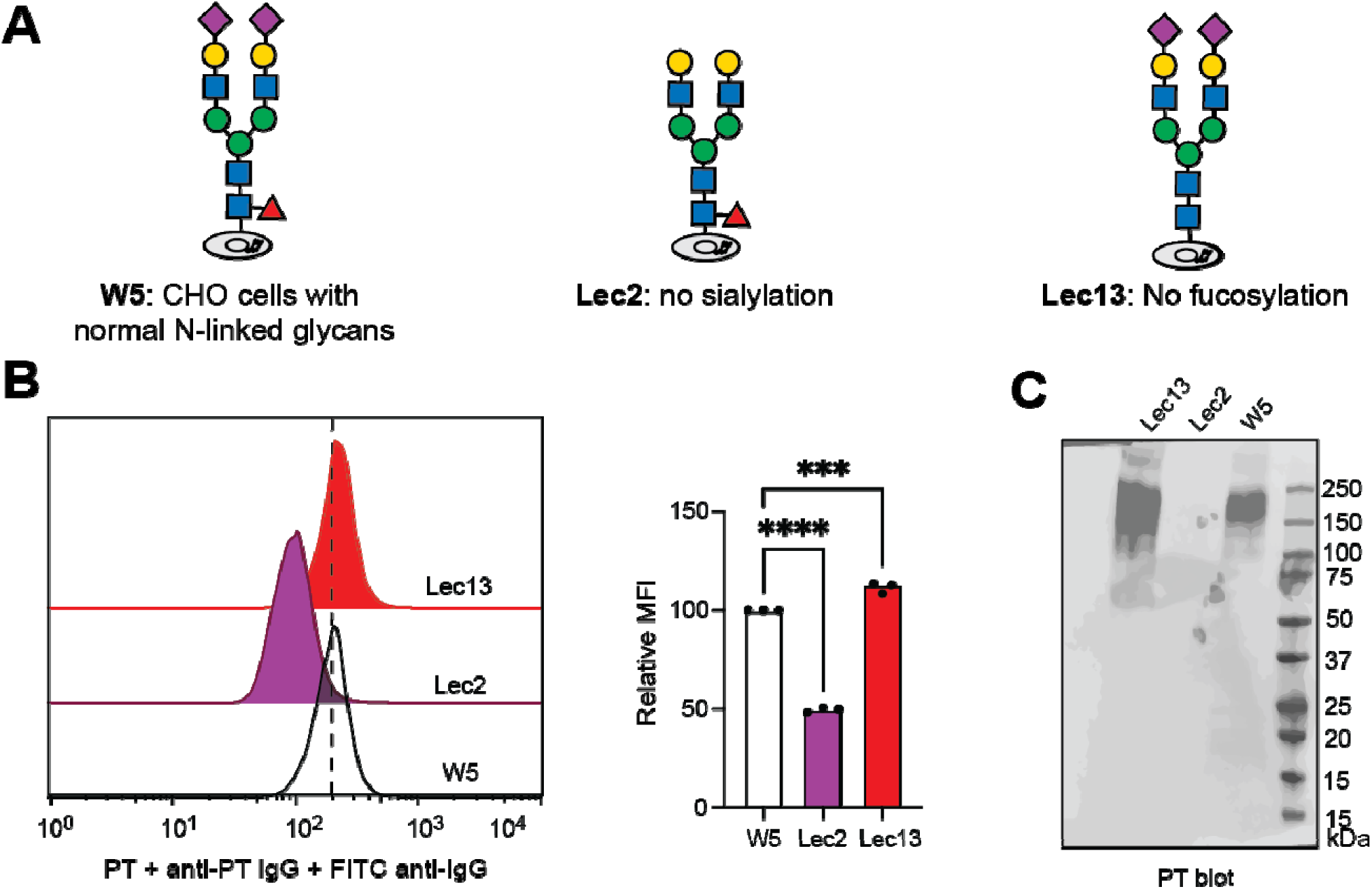
CHO mutant cells lacking fucose on their cell surfaces exhibit increased PT binding. (A). Schematic of mutant CHO cells used in analyzing PT binding. W5: parental CHO cell line; Lec2: mutant CHO cells lacking sialoglycoconjugates; Lec13: mutant CHO cells lacking fucosylated glycoconjugates. **(B**) Flow cytometry analysis of PT binding to cell surfaces of the CHO cells. Bar graph shows the quantification of geometric mean fluorescence from 3 independent trials, normalized to the geometri mean of W5 cells. Statistical analyses were performed by one-way ANOVA (Error bar indicates mean ± SD, *** indicates p < 0.001 and **** p <0.0001). **(C)** PT lectin blot for the lysates of W5, Lec2 and Lec13 CHO cells. The lectin blot shown is representative of 3 biological replicates.

### Lack of fucose results in increased PT binding to Colo205 cells

We recently demonstrated roles for fucosylation in cholera toxin (CT) binding to and intoxication of Colo205 colonic epithelial cells.(51) We took advantage of fucose-deficient Colo205 cells that we used to study CT. CRISPR-Cas9-dependent knockout (KO) of the gene encoding the GDP-fucose transporter (*SLC35C1*) results in global reduction of fucosylation of cell surface glycoconjugates.(51,52) We compared binding of PT to wild-type (WT) and SLC35C1 KO Colo205 cells, as well as non-targeting control (NTCTL) Colo205 cells which express the Cas9 protein and a non-targeting scramble control guide RNA (sgRNA).(51) Flow cytometry revealed greater PT binding to SLC35C1 KO Colo205 cell surfaces compared to WT or NTCTL cells (Fig. 3A). Similarly, lectin blot analysis showed greater PT binding to glycoconjugates from SLC35C1 KO Colo205 cells compared to WT or NTCTL cells (Fig. 3B).

**Figure 3.**
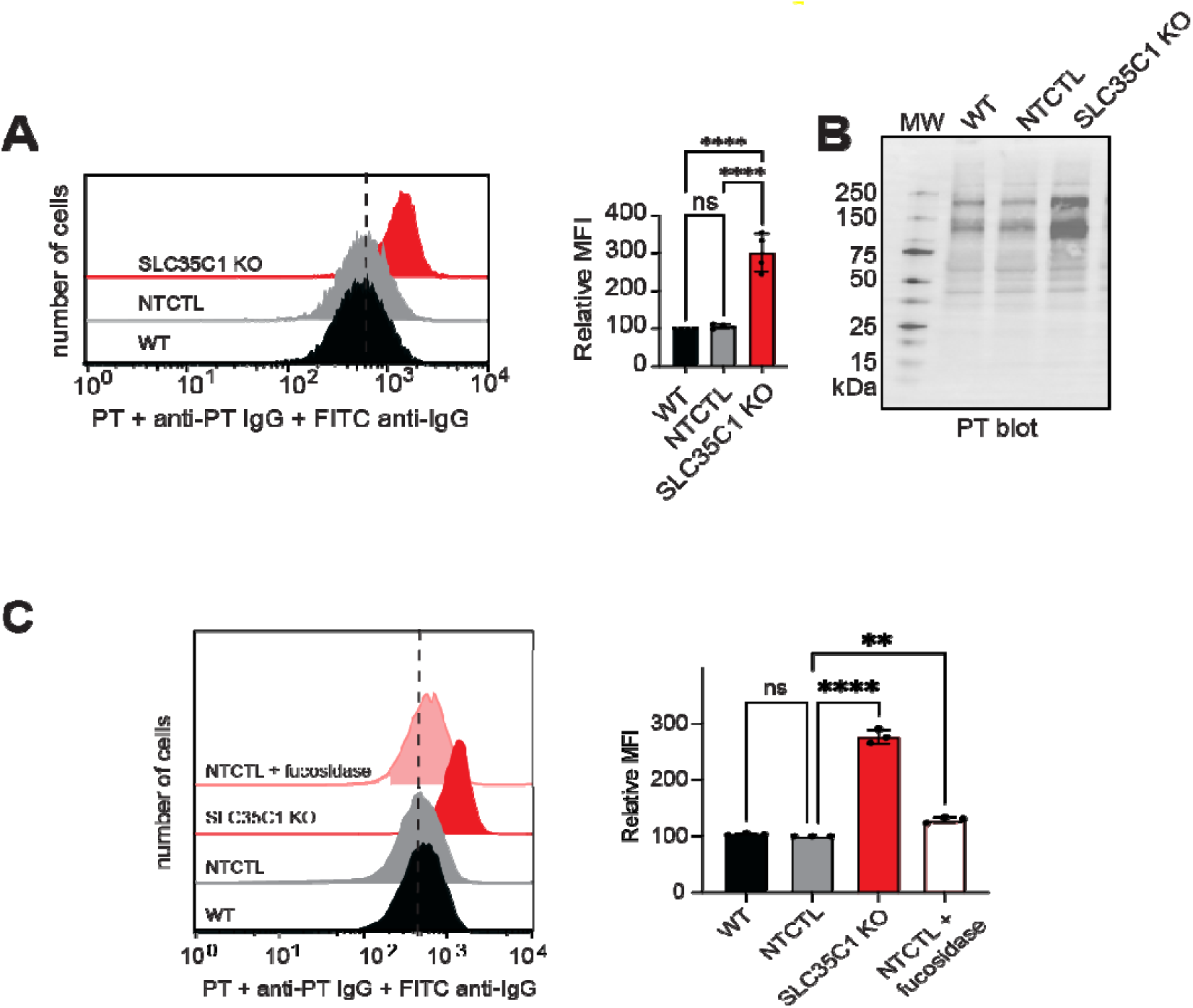
Reduction of fucosylation Colo205 cells results in increased PT binding. **(A**) Flow cytometry analysis of PT binding to cell surfaces of the SLC35C1 KO Colo205 cells. Bar graph shows quantification of geometric mean fluorescence from 3 independent trials, normalized to the geometri mean of wildtype Colo205 cells. **(B)** PT lectin blot for the lysates of wildtype (WT), non-targeted control (NTCTL) and SLC35C1 KO Colo205 cells. The lectin blot shown is representative of 3 biological replicates. **(C)** Flow cytometry analysis of PT binding to cell surfaces of fucosidase-treated Colo205 cells. Bar graph shows the quantification of geometric mean fluorescence from 3 independent trials, normalized to the geometric mean of wildtype Colo205 cells without enzyme treatment. Statistical analyses were performed by one-way ANOVA (error bar indicates mean ± SD, ** indicates p < 0.01; ****p < 0.0001; ns p > 0.05).

One possible explanation for these results is that cells that lack fucosylation produce higher levels of sialylation, a phenomenon observed in HepG2 cells.(53) To assess this possibility, we analyzed the amount of sialic acids on WT, NTCTL, and SLC35C1 KO Colo205 cells using periodate oxidation and aniline-catalyzed oxime ligation (PAL) labeling.(54) This method employs mild periodate oxidation to selectively introduce aldehyde groups on sialic acid residues. The aldehyde groups can be chemoselectively reacted with aminoxy-biotin in the presence of aniline catalyst to biotinylate sialylated glycoconjugates. Streptavidin binding to PAL-labeled WT, NTCTL, and SLC35C1 KO Colo205 cells was analyzed by flow cytometry (Supplementary Fig. 2A). Similarly, the PAL-labeled lysates were analyzed by streptavidin blot (Supplementary Fig. 2B). Neuraminidase treatment showed that biotin labeling was dependent on sialic acid. No measurable difference in biotin labeling was observed between SLC35C1 KO cells and either the WT or NTCTL control cells, suggesting that the higher level of PT binding to SLC35C1 KO cells was not due to a higher level of sialylation.

We also considered the possibility that the pharmacological and genetic interventions to block fucosylation had unanticipated impacts on glycoconjugate biosynthesis or cell surface display of glycoconjugates. To exclude such effects, we used extracellular fucosidase treatment to remove fucose from Colo205 cell surfaces. FucosExo is a mixture of α-fucosidases that removes α1-2, α1-3 and α1-4-linked fucose but not α1-6-linked core fucose. Treatment of Colo205 cell surfaces with FucosExo decreased but did not eliminate cell surface Lewis X (Supplementary Fig. 3A). Colo205 cells treated with FucosExo showed more PT binding than untreated cells, however the magnitude of the increase was smaller than that observed for the SLC35C1 KO (Fig. 3C). This result indicates that at least some of the observed increase in PT binding is due to higher affinity for non-fucosylated glycans as compared to fucosylated glycans. The difference in PT binding observed for SLC35C1 KO cells compared to FucosExo-treated WT cells could mean that the SLC35C1 KO has ancillary effects on glycan structure. Alternately, the difference could be due to incomplete removal of fucose by FucosExo, which does not act on core α1-6-linked fucose and does not completely remove other forms of fucose, including Lewis X.

### Loss of sialyl Lewis X or core fucose is associated with increased PT binding to Colo205 cells

The human genome encodes thirteen fucosyltransferase (FUTs) responsible for adding fucose in specific linkages.(55) To identify the specific forms of fucose that modulate PT binding, we used CRISPR/Cas9 and guide RNAs targeting specific *FUT* genes to prepare Colo205 knockout cell lines. Using a guide RNA targeting *FUT3*, we isolated a monoclonal cell line that displays near-complete loss of sialyl Lewis X expression (FUT3/5/6 3KO.1; Supplementary Fig. 4A). Sequence analysis of genomic DNA from these cells revealed frameshift mutations in all copies of *FUT3*. Additionally, all copies of *FUT5* and *FUT6* contained either frameshift mutations or deletions of two amino acids in the putative donor binding region (Supplementary Fig. 4B). Similarly, using a guide RNA targeting *FUT5*, we isolated an additional monoclonal cell line with near-complete loss of sialyl Lewis X expression (FUT3/5/6 3KO.2; Supplementary Fig. 5A). Sequence analysis of genomic DNA from these cells also revealed frameshift mutations in all *FUT3*, *FUT5*, and *FUT6* genes (Supplementary Fig. 5B). We observed increased PT binding to these two FUT3/FUT5/FUT6 triple KO (3KO) cell lines as compared to WT Colo205 cells (Fig. 4A and 4C). Overexpressing FUT3 or FUT5 in a FUT3/FUT5/FUT6 3KO cell line partially reversed this effect (Fig. 4B and 4D). Additionally, using a guide RNA targeting *FUT8*, we isolated a monoclonal cell line that lacks expression of α1-6-linked core fucose and harbors frameshift mutations in all *FUT8* genes (Supplementary Fig. 6). The FUT8 KO cell line also displayed an increase in PT binding (Fig. 4E), which was reversed by overexpression of FUT8 (Fig. 4F).

**Figure 4.**
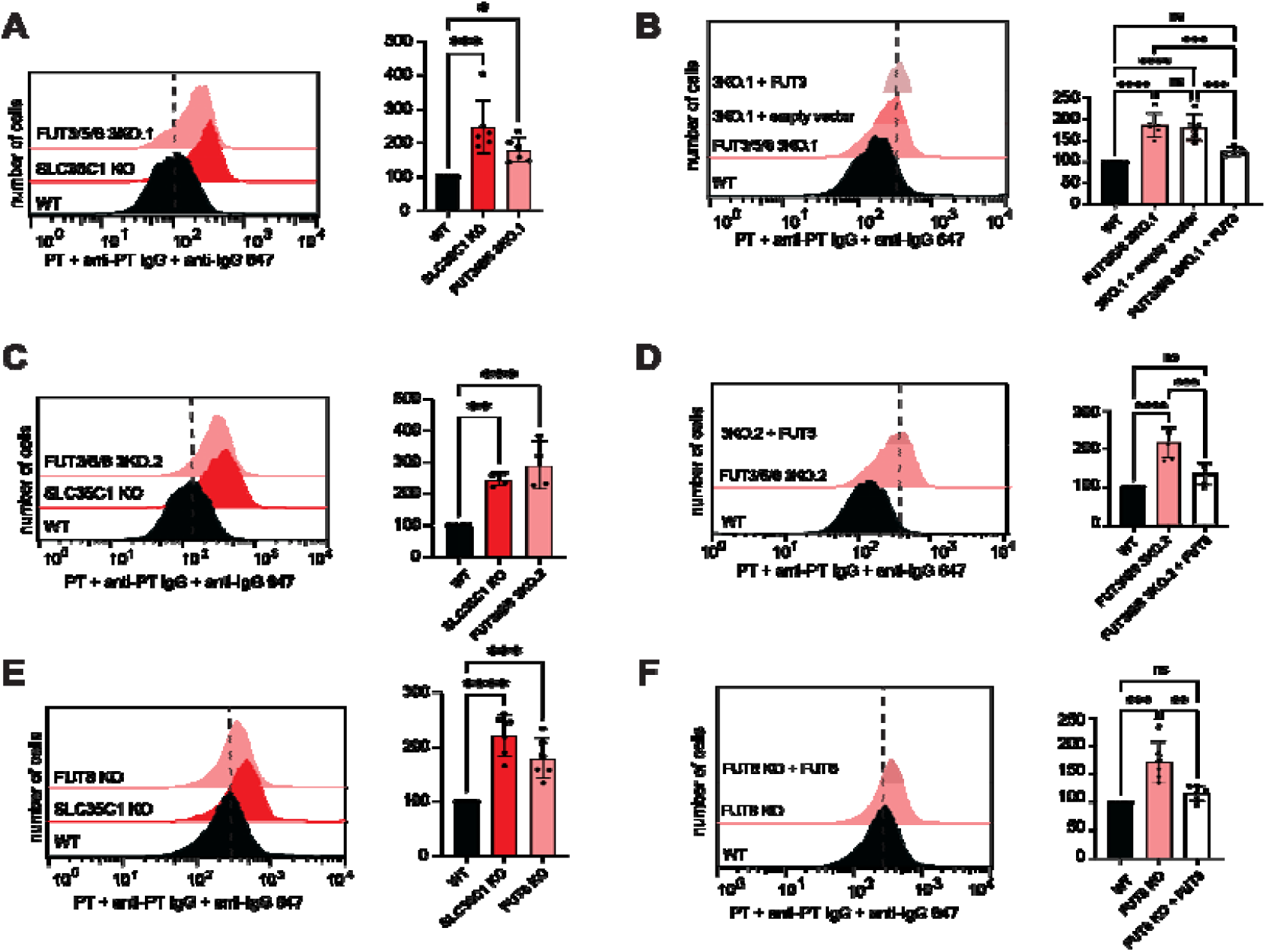
Sialyl Lewis X and core α1-6-linked fucose are each associated with reduced PT binding. **(A)** Flow cytometry analysis of PT binding to cell surfaces of the FUT3/5/6 triple knock out Colo205 cells generated using a guide RNA targeting *FUT3*. Bar graph shows the quantification of geometric mean fluorescence from 6 independent trials, normalized to the geometric mean of WT Colo205 cells. **(B)** Flow cytometry analysis of PT binding to cell surfaces of the FUT3/5/6 triple knock out Colo 205 cells overexpressing FUT3. Bar graph shows the quantification of geometric mean fluorescence from 6 independent trials, normalized to the geometric mean of WT Colo205 cells. **(C)** Flow cytometry analysis of PT binding to cell surfaces of the FUT3/5/6 triple knockout Colo205 cells generated using a guide RNA targeting *FUT5*. Bar graph shows the quantification of geometric mean fluorescence from 4 independent trials, normalized to the geometric mean of WT Colo205 cells. **(D)** Flow cytometry analysis of PT binding to cell surfaces of the FUT3/5/6 triple knockout Colo205 cells overexpressing FUT5. Bar graph shows the quantification of geometric mean fluorescence from 6 independent trials, normalized to the geometri mean of WT Colo205 cells. **(E)** Flow cytometry analysis of PT binding to cell surfaces of the FUT8 knockout Colo205 cells. Bar graph shows the quantification of geometric mean fluorescence from 6 independent trials, normalized to the geometric mean of WT Colo205 cells. **(F)** Flow cytometry analysis of PT binding to cell surfaces of the FUT8 knock out Colo 205 cells overexpressing FUT8. Bar graph shows the quantification of geometric mean fluorescence from 6 independent trials, normalized to the geometri mean of WT Colo 205 cells. Statistical analyses were performed by one-way ANOVA (Error bar indicate mean ± SD, * indicates p < 0.05; **p < 0.01; ***p < 0.001; ****p < 0.0001; ns p > 0.05).

### Reduction of fucose on Jurkat cell surfaces results in increased PT binding and increased ERK phosphorylation

PT is a T cell mitogen.(56) In Jurkat cells, PT promotes signaling through the T-cell receptor (TCR), leading to activation of intracellular signaling cascades including phosphorylation of ERK1 (also known as MAPK3) and ERK2 (also known as MAPK1).(57) To elucidate the impact of fucose on PT function, we used CRISPR/Cas9 to construct Jurkat cells with genetic KO of *SLC35C1* or *FUT8*. Similar to the results from Colo205 cells, both Jurkat KO cell lines displayed increased PT binding as compared to WT or NTCTL cells (Fig. 5A). We then measured PT-induced ERK1/ERK2 phosphorylation. When treated with PT for 1 hour, ERK1/2 phosphorylation was induced in WT cells but not in the SLC35C1 and FUT8 KO cell lines (Fig. 5B). This difference was not due to a loss of ERK pathway components because TPA (12-O-tetradecanoylphorbol-13-acetate) treatment induced similar levels of ERK1/ERK2 phosphorylation in all cell lines (Supplementary Fig. 7A). Rather, this result appears to be consistent with reports that core α1-6-fucosylation is required for signaling through the T-cell receptor.(58–60)

**Figure 5.**
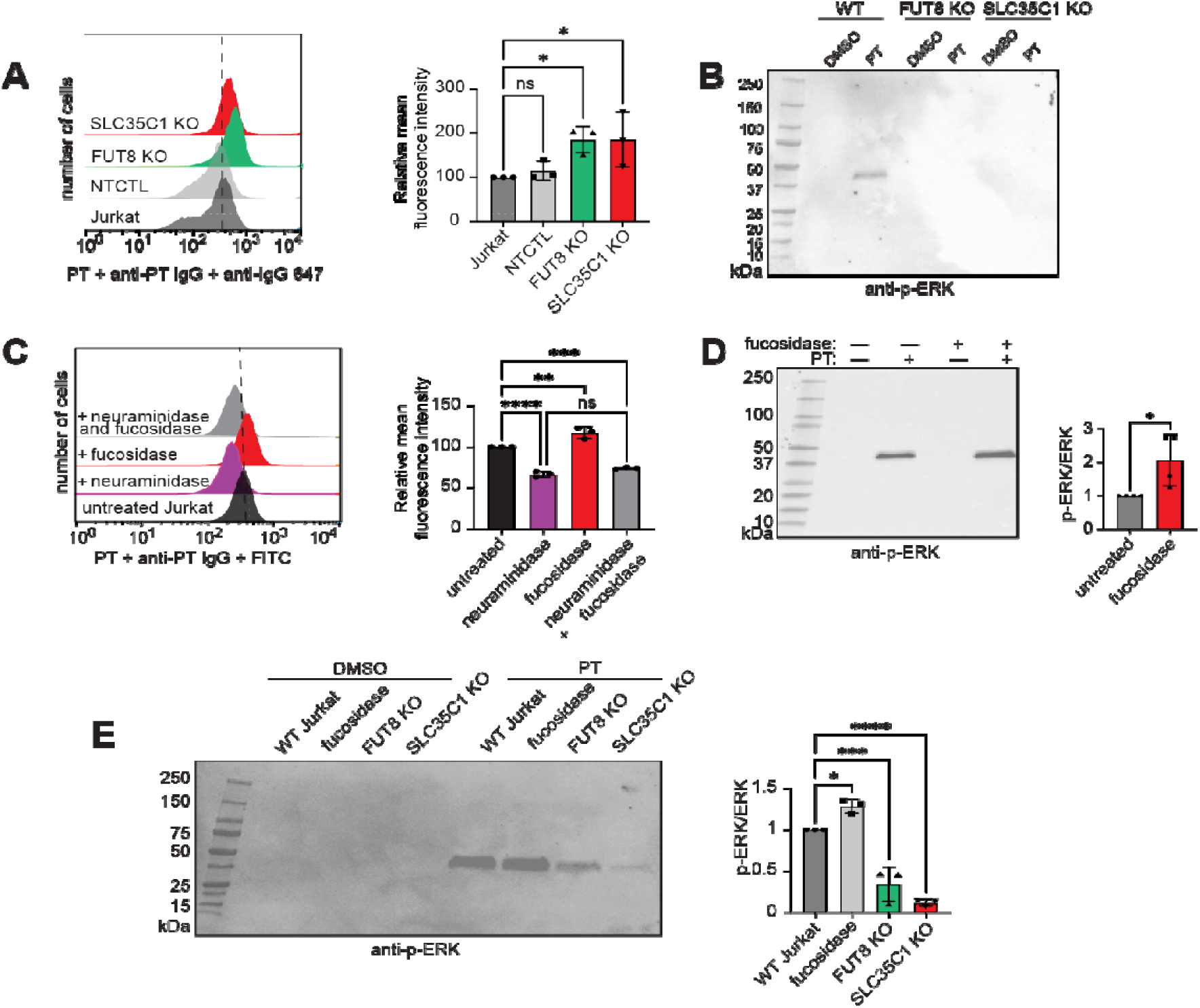
Removal of fucose from Jurkat cell surface glycoconjugates results in increased PT binding and ERK phosphorylation. **(A)** Flow cytometry analysis of PT binding to Jurkat cell surfaces. Bar graph shows the quantification of geometric mean fluorescence from 3 independent trials, normalized to the geometric mean of WT Jurkat cells. **(B)** Phospho-ERK immunoblot of lysates from PT-treated (60 min) Jurkat cells. The blot shown is representative of 3 biological replicates. **(C)** Flow cytometry analysis of PT binding to fucosidase- and neuraminidase-treated Jurkat cell surfaces. Bar graph shows the quantification of geometric mean fluorescence from 3 independent trials, normalized to the geometri mean of WT Jurkat cells. **(D)** Phospho-ERK immunoblot of lysates from PT-treated (60 min) Jurkat cells. The immunoblot shown is representative of 3 biological replicates. **(E)** Phospho-ERK immunoblot of lysates from PT-treated (10 min) Jurkat cells. The immunoblot shown is representative of 3 biological replicates. Statistical analyses were performed by one-way ANOVA (Error bar indicates mean ± SD, * indicates p < 0.05; **p < 0.01; ***p < 0.001; ****p < 0.0001; ns p > 0.05).

To interrogate the role of non-core fucose, we treated wild-type Jurkat cells with FucosExo to remove cell surface α1-2, α1-3, and α1-4-linked fucose but not α1-6-linked core fucose. As a control, we also used *Arthrobacter ureafaciens* neuraminidase to remove α2-3, α2-6, and α2-8-linked sialic acid from Jurkat cell surfaces. As expected, neuraminidase treatment resulted in reduced PT binding, as measured by flow cytometry. In contrast, fucosidase-treated cells exhibited increased PT binding (Fig. 5C), consistent with our observations in other cell types. PT treatment of Jurkat cells for 1 hour led to induced ERK1/ERK2 phosphorylation, as detected by immunoblot (Fig. 5D). This effect was reduced in cells that were pre-treated with neuraminidase (Supplementary Fig. 7B). However, PT-dependent ERK1/ERK2 phosphorylation was enhanced in cells that were pre-treated with fucosidase (Fig. 5D) with no change in total ERK1/ERK2 levels (Supplementary Fig. 7C). Notably, fucosidase treatment alone had no impact on ERK phosphorylation.

Time-course analysis revealed maximal PT-dependent ERK1/ERK2 phosphorylation ten minutes after PT treatment (Supplementary Fig. 7D). Based on this observation, we repeated evaluation of PT-induced ERK1/ERK2 phosphorylation in fucosidase-treated Jurkat cells, as well as FUT8 KO and SLC35C1 KO Jurkat cells, but this time treated with PT for only 10 min. Similar to the experiment performed with 1 hour PT treatment, fucosidase treatment resulted increased sensitivity to PT, resulting in greater ERK1/ERK2 phosphorylation but no change in total ERK1/ERK2 (Fig. 5E and Supplementary Fig. 7E). FUT8 KO and SLC35C1 KO cells were again less sensitive to PT than WT cells but a small amount of ERK1/ERK2 phosphorylation could be observed, suggesting that some residual T-cell receptor signaling can occur in the absence of core fucose.

## Discussion

Here we report that PT exhibits enhanced binding to cells that lack cell surface fucose. Use of genetic, enzymatic, or small molecule inhibitor approaches to reduce fucosylation led to increased PT binding to cell surfaces, as measured by flow cytometry, and to glycoproteins, as assessed by PT blot. The association between hypofucosylation and enhanced PT binding was observed in HBEC-3KT, CHO, Colo205, and Jurkat cell lines.

We further identified fucosyltransferases capable of modulating PT binding through genetic KO experiments. Genetic KO of FUT8 in either Colo205 or Jurkat cells led to increased PT binding, suggesting that core α1-6-fucosylation of N-linked glycans is inhibitory toward PT binding. This result is consistent with data from an oligosaccharide capture ELISA assay that compared PT binding to glycans that differed only in the presence or absence of core α1-6-fucose.(36) PT also exhibited enhanced binding to Colo205 cells that lacked the ability to produce sialyl Lewis X due to genetic KO of FUT3, FUT5, and FUT6. Similarly, enzymatic removal of α1-2, α1-3, and α1-4-linked fucose in Colo205 and Jurkat cells resulted in increased PT binding. This result is consistent with glycan array data that showed that PT binds more strongly to a VIM-2 antigen that contains terminal Neu5Ac-LacNAc, as compared to a sialyl Lewis X glycan.(35) Taken together, our data show that at least two distinct forms of fucosylation are inhibitory to PT binding to cell surface glycoconjugates (Fig. 6). With recent evidence pointing to the existence of multiple glycan binding pockets on S2 and S3,(31) it will be of interest to determine the structural details of how fucosylation modulates glycan binding affinity. Importantly, the results presented here establish the modulatory effect of fucosylation on PT binding to intact cells (Fig. 6).

**Figure 6.**
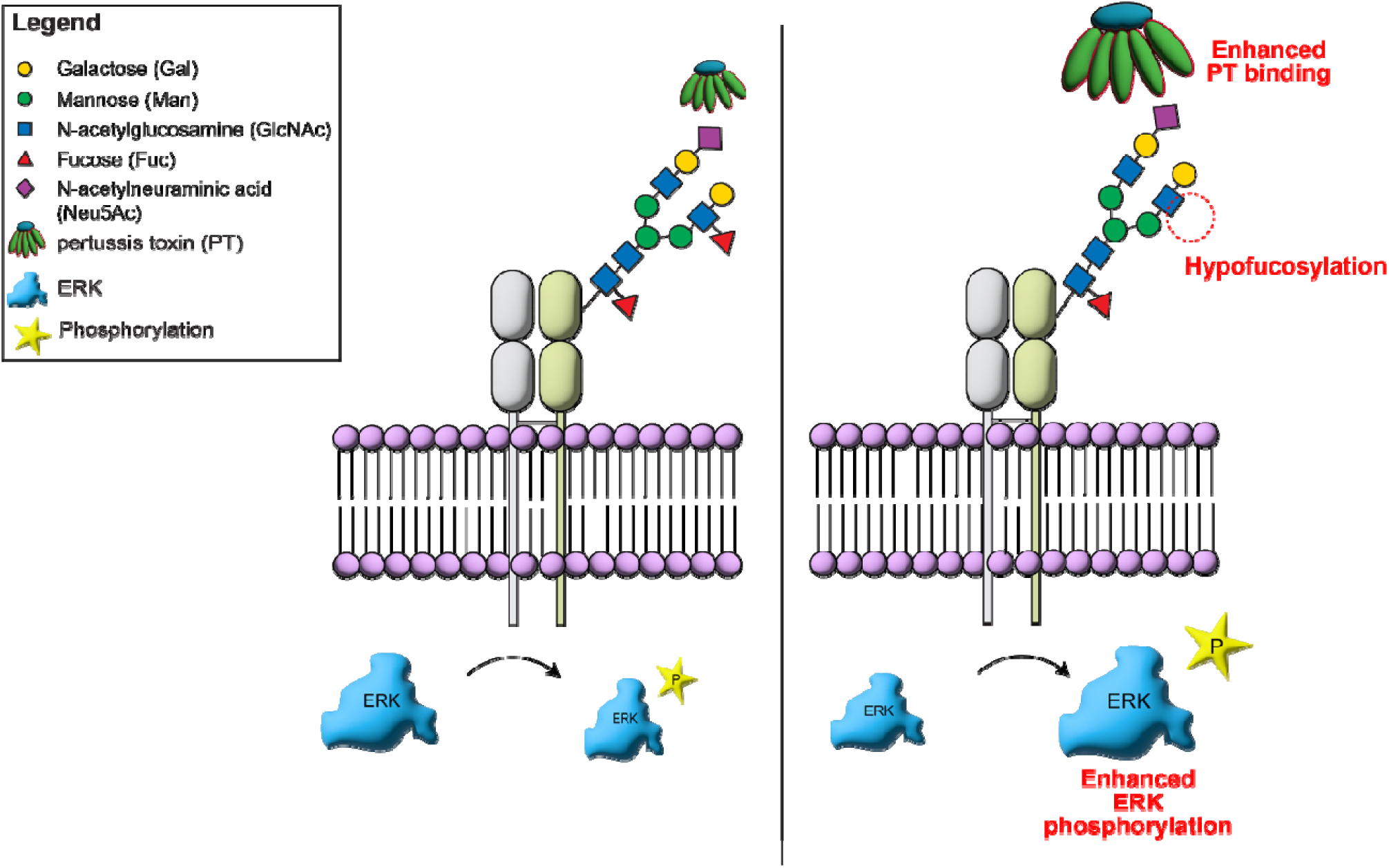
Hypofucosylated glycans enhance PT binding and ERK phosphorylation. PT binds sialylated cell surface glycoconjugates, resulting in intracellular ERK phosphorylation (left). In the hypofucosylated condition (right), binding of PT is increased and results in enhanced phosphorylation of ERK.

We assessed the functional significance of increased PT binding to hypofucosylated glycans by measuring the impact on T cell mitogenicity. Binding of PT activates TCR signaling in Jurkat cells leading to ERK phosphorylation.(57) Despite exhibiting increased PT binding, SLC35C1 and FUT8 KO Jurkat cells showed reduced PT-induced phosphorylation, likely due to the requirement of α1-6 core fucosylation for TCR activity.(58–60) In contrast, Jurkat cells treated with fucosidase to remove α1-2, α1-3, and α1-4-linked fucose exhibited both increased PT binding and increased PT-induced ERK phosphorylation (Fig. 6). Thus, while hypofucosylation generally leads to increased PT binding, the resulting impact on PT activity is dependent on fucose linkage.

One limitation of our study is that all of the tools we used - genetic, enzymatic, or small molecule inhibitor approaches – cause global changes in fucosylation, which can impact not only PT binding but also other cellular processes. Unfortunately, it is not currently feasible to control fucosylation of individual cell surface glycoproteins nor do we know the specific glycoproteins that serve as receptors for PT. Additional work is required to characterize PT receptors from target cell types and this is one of our long term goals. Additionally, we only examined the impact of hypofucosylation on PT mitogenicity. PT exhibits other cellular activities, including the ability to intoxicate cells through internalization and ADP-ribosylation of Gαi proteins. Future efforts will focus on examining the impact of fucosylation on other PT functions.

Our study demonstrates the role of fucose in the modulation of PT binding and function. This result implies that physiological states that exhibit altered fucosylation of PT target cells could lead to changes in disease severity. For example, recent studies have demonstrated increases in fucosylation of airway epithelial cells in inflammatory states induced by allergens or mechanical injury.(61–63) Such changes in glycosylation could serve to protect the host against PT. Currently, the primary treatments for pertussis are antibiotics, which are only effective in the early stage of infection. Treatments that target PT are of potential interest to treat later stages of disease.(64) Understanding the molecular mechanisms of PT action could facilitate development of therapies that target PT binding to host cells, such as competitive inhibitors of PT binding or treatments that modulate host cell glycosylation.

## MATERIALS AND METHODS

### Chemicals

Glycan synthesis inhibitors peracetylated 3-fluoro-β-D-N-acetylneuraminic acid (3F_ax_Neu5Ac) (cat. # 566224), peracetylated 2-deoxy-2-fluoro-L-fucose (2F-Fuc) (cat. # 344827), and kifunensine (cat. # 422500) were purchased from Sigma. Stock concentrations were made in dimethyl sulfoxide (DMSO; cat. # D8418, Sigma). Bovine serum albumin (BSA) and skim milk powder (cat. # NC9121673) were purchased from Fisher Scientific. Carbohydrate-free blocking solution was purchased from Vector Laboratories (Burlingame, CA) (cat. # SP-5040). COmplete ULTRA protease inhibitor tablets were purchased from Roche.

### Pertussis toxin

Pertussis toxin (PT) was purchased from List Labs (cat. # 180). The lyophilized toxin was dissolved in 200 µL Dulbecco’s Phosphate Buffered Saline (DPBS) (cat. # D8537, Sigma) to achieve a stock concentration of 200 µg/mL. The resuspended toxin was stored at 4 °C until use.

### Cell culture

All cell lines used in the current study were cultured at 37 °C, 5 % CO_2_ in a water-saturated environment. HBEC-3KT cells were obtained from John Minna (UT Southwestern Medical Center).(65) Colo205 and Jurkat cells were obtained from the ATCC. W5, Lec2, and Lec13 CHO cells were obtained from Pamela Stanley (Albert Einstein College of Medicine). HBEC-3KT cells were grown in 1 % porcine gelatin cell culture dishes containing Keratinocyte-SFM (ThermoFisher Scientific cat. # 17005-042) growth medium reconstituted with human recombinant epidermal growth factor (EGF 1-53) and bovine pituitary extract (BPE). CHO cells were cultured in alpha MEM medium (GIBCO 11900-073) supplemented with 10 % fetal bovine serum (FBS). Lec2 and Lec13 cells were cultured in media containing dialyzed FBS (cat # A3382001, GIBCO) to minimize availability of free sialic acid and fucose. Colo205 and Jurkat cells were cultured in RPMI 1640 medium (cat #11875093, GIBCO) supplemented with 10 % FBS, and 1 % penicillin/streptomycin (cat #15140122 , GIBCO).

### Flow cytometry

For flow cytometry experiments, 5 x 10^5^ cells were resuspended in 100 µL DPBS containing 1 % (w/v) BSA (flow buffer) and were added per well to a V-bottom plate (Costar cat. # 3897). For analyzing PT binding, resuspended cells were incubated for 40 min on ice with 100 µL of 1 μg/mL of PT. Cells were washed twice with 200 μL of cold flow buffer and incubated with 100 μL of 1:1000 dilution of anti-beta subunit PT (Abcam, cat. # ab188414). For detection of fucosylated glycoconjugates and Lewis X, cells were incubated for 30 minutes on ice with 50 μL of either 1:500 dilution of biotinylated *Aleuria aurantia* lectin (AAL, Vector labs cat # B-1395-1) or 1:500 dilution of mouse anti-human CD15 clone HI98 (BD Biosciences, cat. # 555400BD). For the detection of sialylated glycans, 50 µl of 2.5 µg/mL biotinylated Pan-Siafind (Lectenz, cat # SK0501) was used. Cells were washed twice with flow buffer and incubated with 50 μL of 1:500 dilution of respective secondary reagents for 30 min on ice. For PT, the secondary reagent was goat anti-rabbit IgG Alexa Fluor® 647 (Abcam, cat # ab150079) or goat anti-rabbit FITC (Abcam, cat # ab ab6717); for Le^x^, the secondary antibody was goat anti-mouse IgM Alexa Fluor® 647 (Abcam, ab150123); for AAL and Pan-Siafind, the secondary reagent was streptavidin-APC (Invitrogen, cat # SA1005). Cells were washed three times with 200 μL of flow buffer and then resuspended in 500 μL flow buffer containing 1 μg/mL propidium iodide. Samples were analyzed on a FACS Calibur flow cytometer (BD Biosciences, UTSW core facility). Cells were gated based on forward versus side scatter, and dead cells were excluded based on PI staining on FL3 channel. For each sample, 10,000 live cell events were measured, and the fluorescence intensity was determined on either FL1 or FL4 channel. Flow cytometry data were analyzed using the FlowJo software.

### Construction of Colo205 KO cell lines

Predesigned CRISPR sgRNAs were ordered from Millipore Sigma and ligated into a neomycin-resistant encoding lentiviral vector, LentiGuide-Neo (Addgene, plasmid no. 139449) as previously described.(66) Ligated plasmids were purified using ZymoPURE^TM^ II Plasmid Midiprep Kit according to manufacturer’s instructions and validated by DNA sequencing.

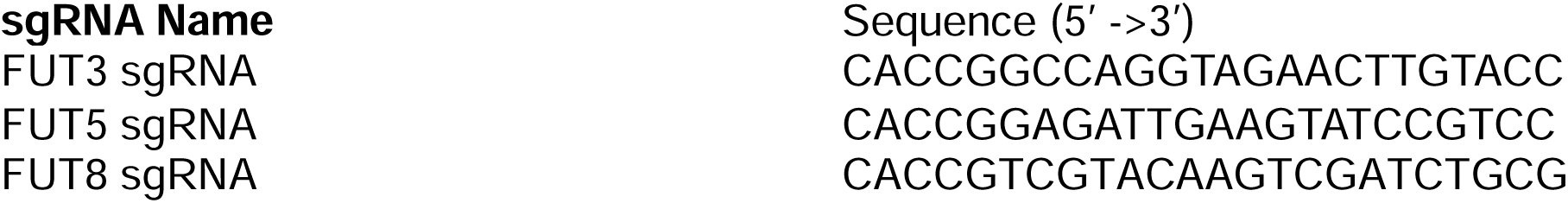

To generate lentivirus, 5×10^5^ low-passage HEK293T\17 cells (ATCC) were plated in individual wells of a 6-well plate in 5 mL DMEM media containing 10% FBS and penicillin/streptomycin. After 24 h,1000 ng of the sgRNA- or Cas9-encoding plasmid, 900 ng of psPax2 packaging plasmid, and 100 ng of pMD2.G packaging plasmid were added to OPTI-MEM media (GIBCO, cat# 31985-062), adjusting the final volume to 1 mL. Room temperature X-tremeGENE 9 DNA Transfection Reagent (6 μL; Sigma Aldrich, LOT33813300) was added to the OPTI-MEM and plasmid mixture. After 15 min, the transfection mixture was added to HEK283T\17 cells in a drop-wise fashion. Cells were incubated at 37 °C in 5% CO_2_ for 24 h, then serum was added to a final concentration of 30%. After 48 h, viral supernatants from cells were collected and filtered through a 0.45 micron filter.

To transduce Colo205 cells, 1×10^5^ cells were plated per well in a 12-well plate containing 2 mL RPMI complete media. Varying volumes of FUT lentivirus and Cas9 lentivirus (50 μL, 100 μL, 200 μL, and 500 μL) were added in equal amounts to RPMI complete media containing 8 μg/mL polybrene for a total final volume of 2 mL. Viral dilutions were added in a dropwise fashion to each well and spinfected at 1000*g* for 2 hours at 33 °C. After spinfection, cells were incubated for 48 hours. Cells were selected with 900 μg/mL G418 and 10 μg/mL blasticidin. To create monoclonal cell lines, polyclonal cells underwent single cell sorting using a BD FACSAria cell sorter at UT Southwestern Flow Cytometry Core facility.

To sequence monoclonal populations, genomic DNA (gDNA) was isolated from each cell line using QuickDNA^TM^ Miniprep Plus Kit (ZymoResearch, cat # D4068) according to the manufacturer’s instructions. sgRNA sequences were amplified by PCR using EmeraldAmp MAX HS PCR Master Mix (Takara Bio, catalog no. RR330B) according to manufacturer’s instructions. The forward and reverse primers for each gene are listed below. PCR products were submitted to Eurofins Genomics for nanopore sequencing and sequence results was analyzed using CRISPResso2 software.(67)

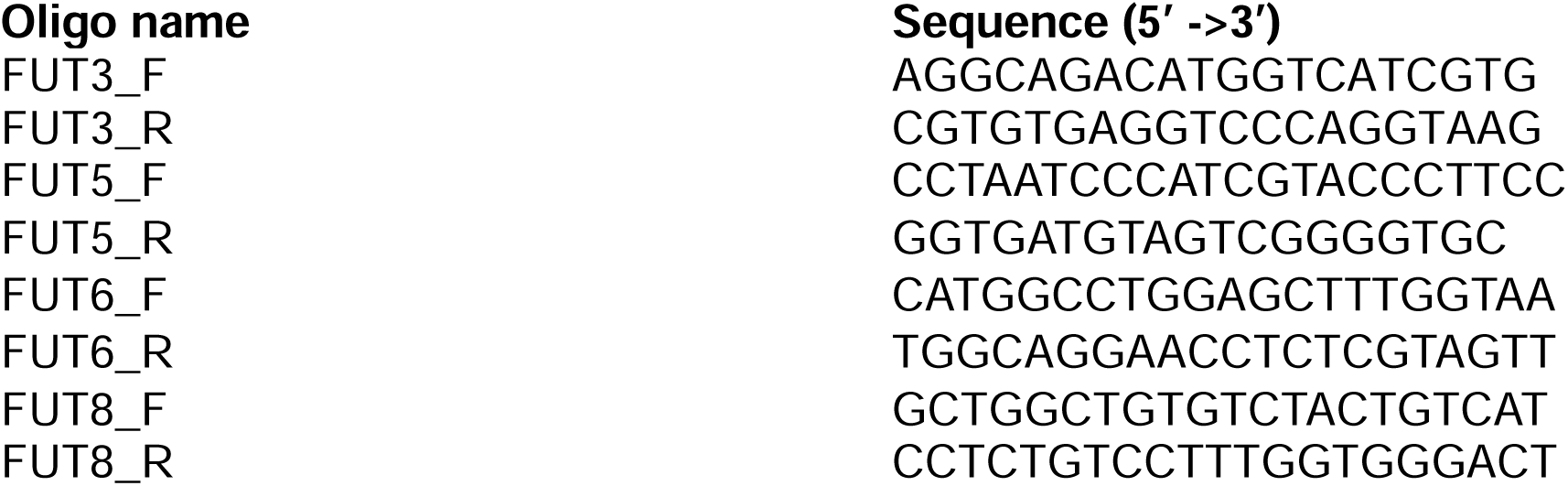

### Construction of Colo205 rescue cell lines

Plasmids encoding the *FUT* open reading frames were obtained from Invitrogen Ultimate ORF clone library (*FUT3*), Genscript (*FUT5*; cat # OHu29705), Origene (*FUT8*; cat # RC223075). Silent mutations to the protospacer-adjacent motif (PAM) site were introduced by mutagenic primers using the QuikChange II XL Site-Directed Mutagenesis Kit (Agilent, cat # 200521). Open reading frames of *FUT3, FUT5 and FUT8* containing mutated PAM sites were amplified by Q5 High Fidelity 2X Master Mix (NEB, cat # M0492L). PCR products were purified by NucleoSpin® Gel and PCR Clean-Up (Takara Bio, cat # 740609.5) according to manufacturer’s instructions. Purified ORFs containing mutated PAM sites were ligated into a pLVX-IRES-puromycin vector (Takara Bio, cat # 632183) linearized by EcoRI-HF and BamHI-HF (NEB, cat # R3101S and R3136S) using HiFi DNA assembly master mix (NEB cat # E5520S). Lentivirus was generated as above and cell lines were spinfected as above. Rescue cell lines were selected with 5 µg/mL puromycin.

### Lectin and immunoblots

Harvested cells (1 x 10^6^), after washing with DPBS buffer, were resuspended and incubated for 30 min on ice in 200 μL RIPA buffer (20 mM Tris-HCl pH 8; 150 mM NaCl; 1 % (v/v) Triton X-100; 0.1 % (v/v) sodium deoxycholate; 1 % (v/v); 1X protease inhibitor). Cell lysates were centrifuged at 14,000*g* for 10 min at 4 °C and the supernatant was transferred to a new microcentrifuge tube.

Total protein concentration was quantified using BCA protein assay kit (cat # 23225, Thermo Fisher Scientific). For PT lectin blot analysis, samples were denatured with Laemmli sample buffer containing 50 mM DTT and were resolved on 4-20 % stain-free gradient SDS-PAGE (cat # 4568094, Bio-Rad Laboratories). Resolved proteins were transferred to PVDF membrane at 100 V for 2 h. Membranes were blocked with 1X Carbo-Free buffer (cat # SP-5040-125, Vector Laboratories) for 1 h at room temperature and then incubated with 2 μg/mL pertussis toxin at 4 °C overnight. The membrane was washed three times using Tris-Buffered Saline with Tween-20 (TBST) and further incubated with 1:1000 rabbit anti-pertussis toxin antibody (cat # Ab188414, Abcam) for 1 h at room temperature. The washing step was repeated with TBST and the membrane was further incubated with 1:2000 goat anti-rabbit IgG-HRP conjugate (Invitrogen cat # 65-6120) for 1 h at room temperature. Washing steps were repeated and the membrane was developed using SuperSignal™ West Pico PLUS Chemiluminescent Substrate (cat #. 34580, Thermo Scientific™) and imaged on a ChemiDocTM MP Imaging System (Bio-Rad). For phosphorylated ERK immunoblots, membranes were blocked in 5 % (w/v) non-fat milk (NFM) in PBST for 1 h at room temperature and then incubated with 1:2000 anti-phospho-p44/42 MAPK (Erk1/2) antibody (cat # 9106, Cell Signaling Technology) overnight at 4 °C in PBST containing 5 % (w/v) NFM. Membranes were washed and then incubated with a goat anti-mouse antibody conjugated to HRP (1:2000, Invitrogen cat # 31430) in PBST containing 5 % (w/v) non-fat milk. For analyzing total ERK immunoblots, membranes were blocked in 5 % (w/v) NFM (non-fat milk) PBST for 1 h at room temperature and then incubated with 1:1000 anti-p44/42 MAPK (Erk1/2) antibody (cat # 4695, Cell Signaling Technology) overnight at 4 °C in PBST containing 5 % (w/v) NFM. Membranes were washed and then incubated with 1:2000 goat anti-rabbit IgG-HRP conjugate in PBST containing 5 % (w/v) NFM. After washing, membranes were developed using SuperSignal™ West Pico PLUS Chemiluminescent Substrate, imaged on a ChemiDocTM MP Imaging System (Bio-Rad).

### PAL (periodate oxidation and aniline-catalyzed oxime ligation) labelling

Colo205 cells (1 x 10^6^) were washed in DPBS and resuspend in 1 mM sodium periodate (Sigma) on ice for 30 min. The oxidation reaction was quenched by adding an equal volume of 1 mM glycerol before washing with PBS. Washed cells were incubated in PBS, containing 5% (w/v) BSA, 250 μM aminooxy-biotin (Biotium), and 10 mM aniline at 4 °C with constant rotation to perform the oxime ligation. Biotin-labeled cells were analyzed using either flow cytometry or immunoblot.

### Glycosidase treatment

Colo205 or Jurkat cells (1 x 10^6^) were washed in DPBS and resuspended in flow buffer. To the resuspended cells, 0.02U of neuraminidase from *Arthobacter ureafaciens* (cat. # 10269611001, Roche) was added and incubated for 1 h with continuous rotation at 37 °C. Enzyme-treated cells were washed with flow buffer and analyzed using flow cytometer or immunoblots.

Cells (1 x 10^6^) were washed in DPBS and resuspend in flow buffer. To the resuspended cells, 80 units of FucosEXO™ (cat. # G1-FM1-020, Genovis) was added and incubated for 1 h with continuous rotation at 37 °C. Enzyme-treated cells were washed with flow buffer and analyzed by flow cytometry or immunoblot.

### ERK phosphorylation analysis

1×10^6^ Jurkat cells were stimulated for 1 h with 2 μg/mL PT or 200 μM TPA (12-O-Tetradecanoylphorbol-13-acetate). After incubation, cells were washed with PBS and lysed with RIPA buffer. Cell lysates were analyzed for phosphorylated ERK 1 and 2 (phospho-ERK) and total ERK by immunoblot, as described above.

## ACKNOWLEDGEMENTS

This work was supported by the Welch Foundation (I-1686) and the NIH (R21AI183574). NN was supported by a postdoctoral fellowship from the German Academic Exchange Service (Deutscher Akademischer Austauschdienst, DAAD). As a part of PB2PHD program, Aurora Silva was supported by The Department of Biochemistry (Chilton Foundation) and the UTSW Graduate School of Biomedical Sciences. We thank Pamela Stanley (Albert Einstein) for sharing CHO cell lines and John Minna (UT Southwestern) for sharing HBEC-3KT cells. We thank Nicholas Carbonetti (University of Maryland School of Medicine) for advice and reagents that were used in pilot studies not described here.

## DATA AVAILABILITY STATEMENT

Datasets and replicate blots will be available in the Texas Data Repository.

LentiGuideFUT3-neo, LentiGuideFUT5-neo, LentiGuideFUT8-neo, pLVX-FUT3-IRES-puro, pLVX-FUT5-IRES-puro, and pLVX-FUT8-IRES-puro plasmids will be available from Addgene.

