## Supplemental Figures for "Hypofucosylation promotes pertussis toxin binding to cell surface glycococonjugates and pertussis toxin-induced intracellular ERK signaling"

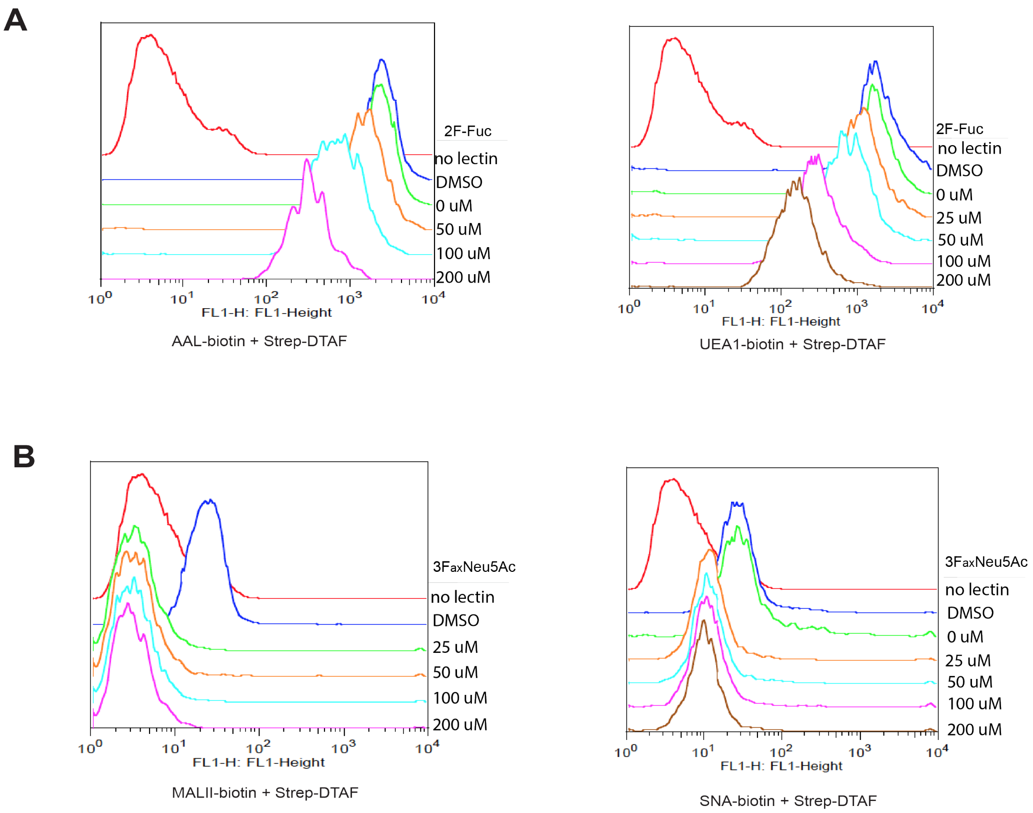


**Supplementary Figure 1. Effectiveness of small molecule inhibitors of glycosylation in HBEC-3KT cells.** HBEC-3KT cells were incubated in media containing a range of glycan inhibitor concentrations to identify the concentration required for successful inhibition of glycan biosynthesis. Cell surface glycans were subsequently analyzed by measuring lectin binding using flow cytometry. **(A)** Flow cytometry histograms showing binding of fucose-specific lectins to HBEC-3KT cell surfaces after culturing with 0, 25, 50, 100, and 200 μM 2-fluorofucose for 72 h. AAL-biotin (left panel) was used to detect multiple forms of fucose and UEA1-biotin (right panel) was used to detect $\alpha$1-2-linked fucose. Streptavidin-DTAF was used as the secondary detection reagent. **(B)** Flow cytometry histograms showing binding of sialic acid-specific lectins to HBEC-3KT cell surfaces after cultured cells with 0, 25, 50, 100, and 200 μM 3-FaxNeuAc for 72 h. MAL II-biotin (left panel) was used to detect $\alpha$2-3-linked sialic acid and SNA-biotin (right panel) was used to detect $\alpha$2-6-linked sialic acid. Streptavidin-DTAF was used as the secondary detection reagent.

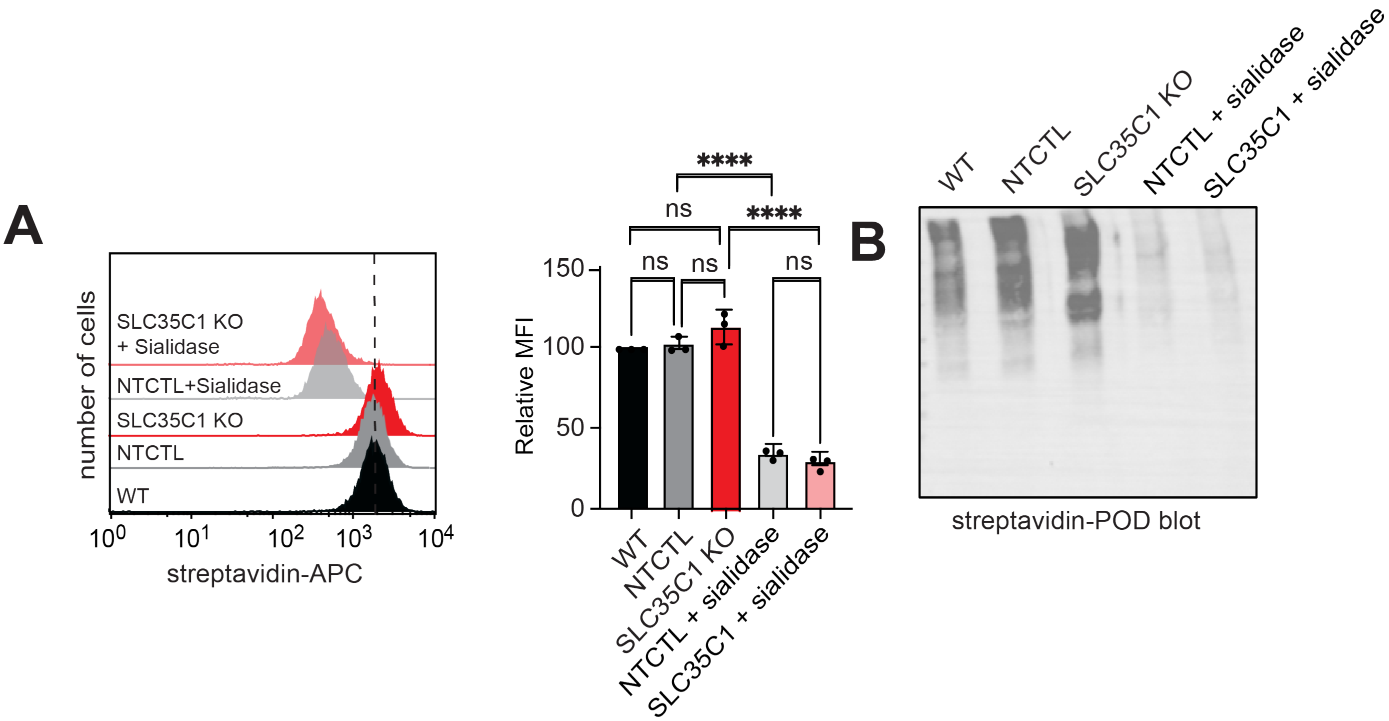


**Supplementary Figure 2. Impact of SLC35C1 KO on sialylation.** Colo205 cell surface sialic acid was biotinylated by PAL labelling. **(A)** Flow cytometry analysis of PAL-dependent biotinylation of Colo205 cell surfaces. Bar graph shows the quantification of geometric mean fluorescence from 3 independent trials, normalized to the geometric mean of wild type Colo205 cells. **(B)** Streptavidin-POD (peroxidase) blot of lysates from PAL-biotinylated wild type (WT), non-targeted control (NTCTL), and SLC35C1 KO Colo205 cells. Statistical analyses were performed by one-way ANOVA. Error bar indicates mean ± SD, **** indicates $p < 0.0001$; ns $p > 0.05$.

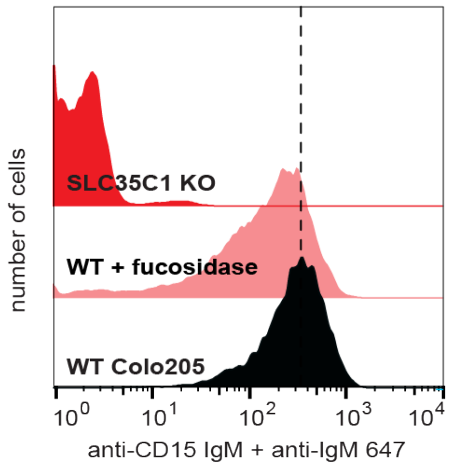


**Supplementary Figure 3. Fucosidase treatment reduces Lewis X levels on the surface of Colo205 cells.** Flow cytometry histograms showing the binding of anti-Lewis X (anti-CD15) antibody to Colo205 cells. WT Colo205 cells were treated with FucoExo™, a mixture of fucosidases, to remove α1-2, α1-3, and α1-4 linked fucose from the cell surface. Fucosidase treatment resulted in a modest reduction in Lewis X as compared to genetic knockout of *SLC35C1*, which eliminated Lewis X expression.

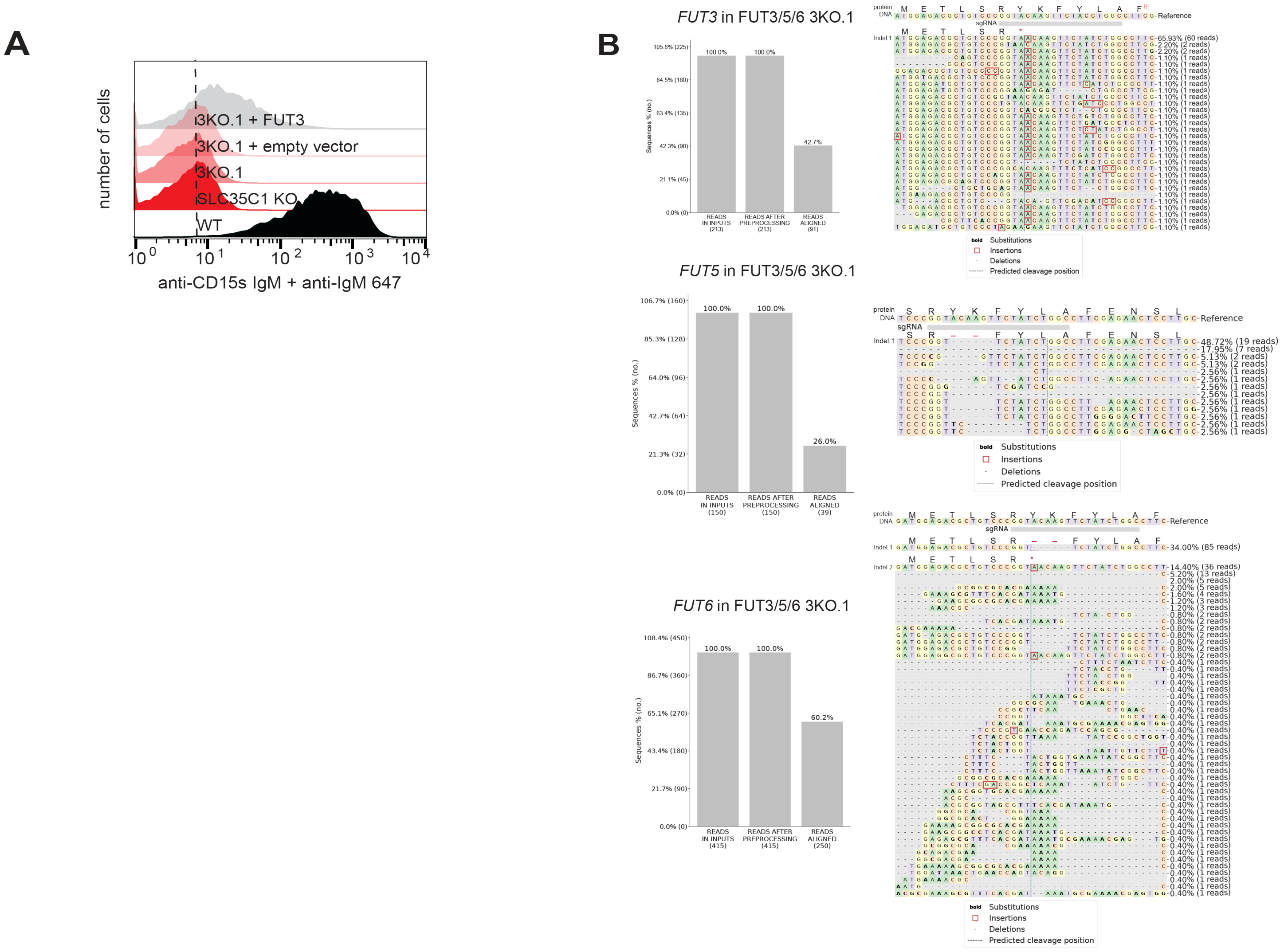


**Supplementary Figure 4. Characterization of FUT3/5/6 TKO.1 Colo205 cells.** (**A**) FUT3/5/6 TKO.1 Colo205 cells were phenotypically characterized by measuring sialyl Lewis X antibody binding. Flow cytometry histogram showing the binding of mouse anti-human sLewis X (BD Pharmingen™ 551344) to Colo205 cells. Anti-mouse IgM-647 was used as secondary detection reagent. (**B**) *FUT3*, *FUT5*, and *FUT6* loci from FUT3/5/6 TKO.1 Colo205 cells were amplified by PCR; the PCR product was sequenced with Oxford nanopore technology. Sequence data were analyzed with CRISPResso2 software. Bar graphs represent the total number of reads and aligned reads. Allele frequency table shows mutations introduced to individual reads. The reference sequence for each cell line shows where the sgRNA binds in the genome. The translated protein sequence is provided for sequences that were detected with more than 15 reads. Detected indels result in a frameshift mutations or two amino acid deletions within the donor-binding domain.

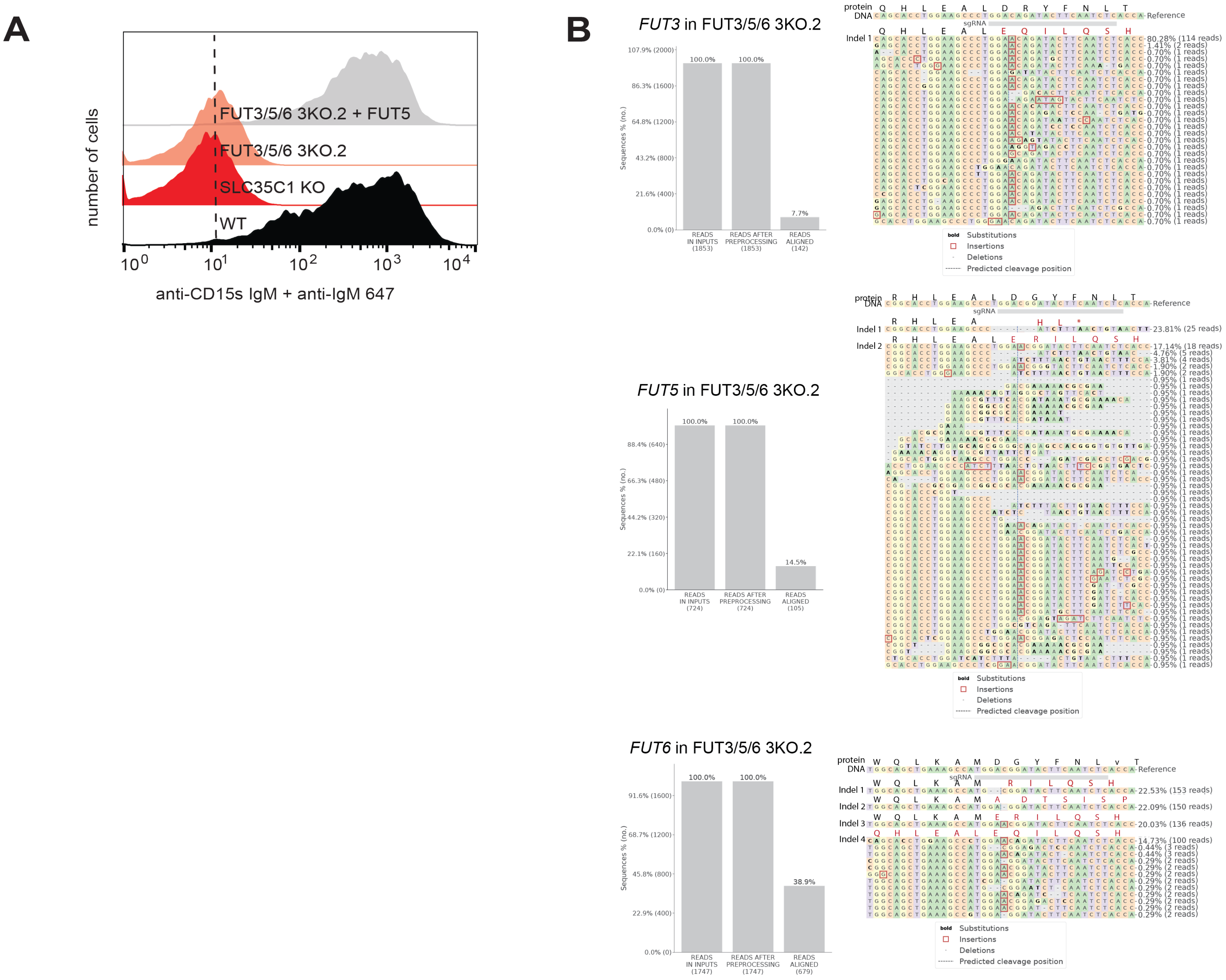


**Supplementary Figure 5. Characterization of FUT3/5/6 TKO.2 Colo205 cells. (A)** FUT3/5/6 TKO.2 Colo205 cells were phenotypically characterized by measuring sialyl Lewis X antibody binding. Flow cytometry histogram showing the binding of mouse anti-human sLewis X (BD Pharmingen™ 551344) to Colo 205 cells. Anti-mouse IgM-647 was used as secondary detection reagent. **(B)** *FUT3*, *FUT5*, and *FUT6* loci from FUT3/5/6 TKO.2 Colo205 cells were amplified by PCR; the PCR product was sequenced with Oxford nanopore technology. Sequence data were analyzed with CRISPResso2 software. Bar graphs represent the total number of reads and aligned reads. Allele frequency table shows mutations introduced to individual reads. The reference sequence for each cell line shows where the sgRNA binds in the genome. The translated protein sequence is provided for sequences that were detected with more than 15 reads. All detected indels result in a frameshift mutations.

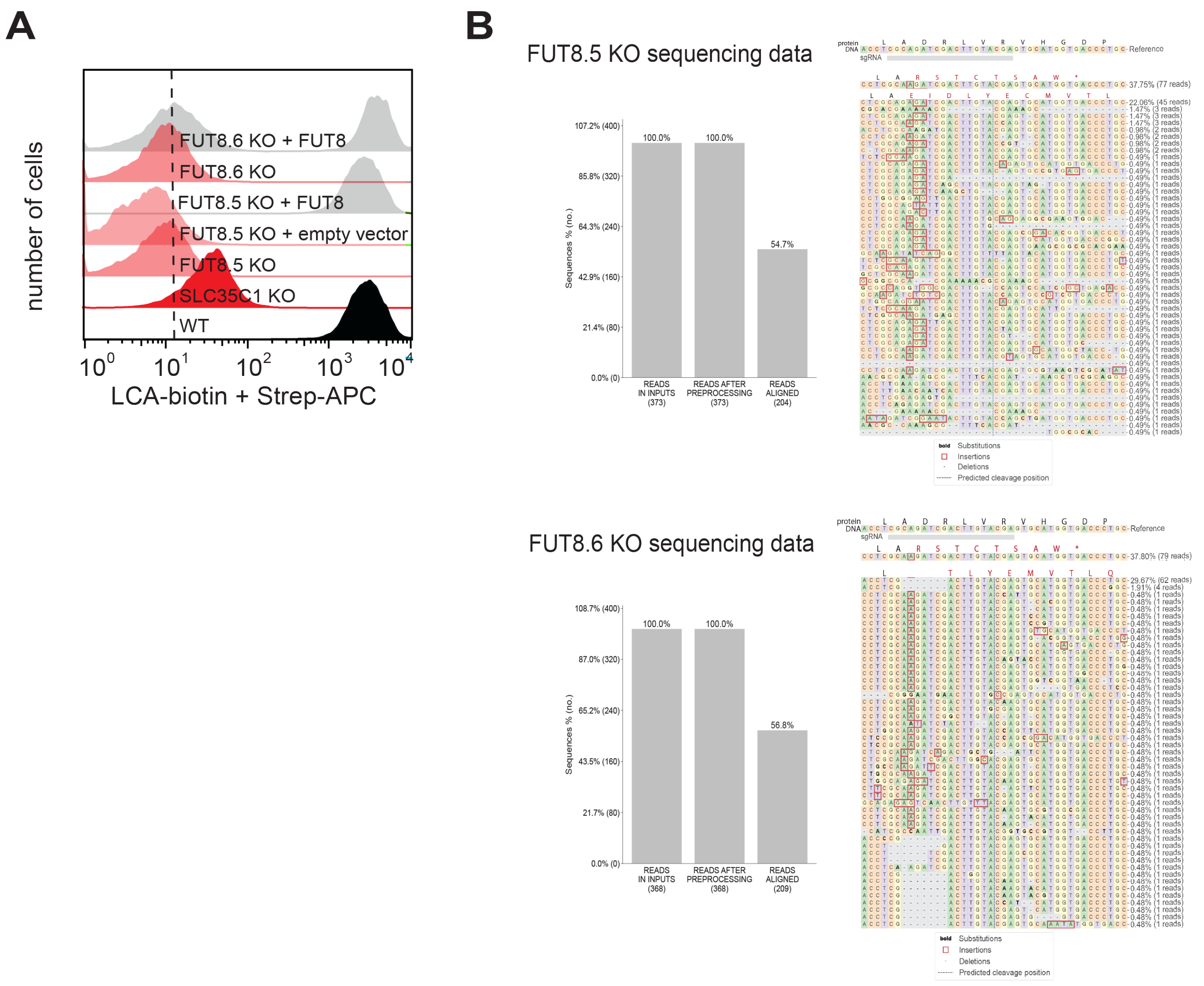


**Supplementary Figure 6. Characterization of FUT8 KO Colo205 cells. (A)** The FUT8 KO Colo205 cells were phenotypically characterized by measuring LCA binding. Flow cytometry histogram showing the binding of LCA-biotin (Vector labs B-1045-5) to Colo205 cells. Streptavidin-APC (allophycocyanin) was used as secondary detection reagent. **(B)** *FUT8* genomic DNA from FUT8 KO Colo205 cell lines were amplified by PCR and the PCR product was sequenced with Oxford nanopore technology. Sequence data were analyzed with CRISPResso2 software. Bar graphs represent the total number of reads and aligned reads. Allele frequency table shows mutations introduced to individual reads. The reference sequence for each cell line shows where the sgRNA binds in the genome. The translated protein sequence is provided for sequences that were detected with more than 15 reads. Both indels for FUT8.5 KO result in frameshift mutations at amino acids 314. Indel 1 for FUT8.6 KO results in a frameshift mutation at amino acid 314, while indel 2 results in a frameshift mutation at amino acid 313. FUT8.5 KO cells were used for all data presented in the main text.

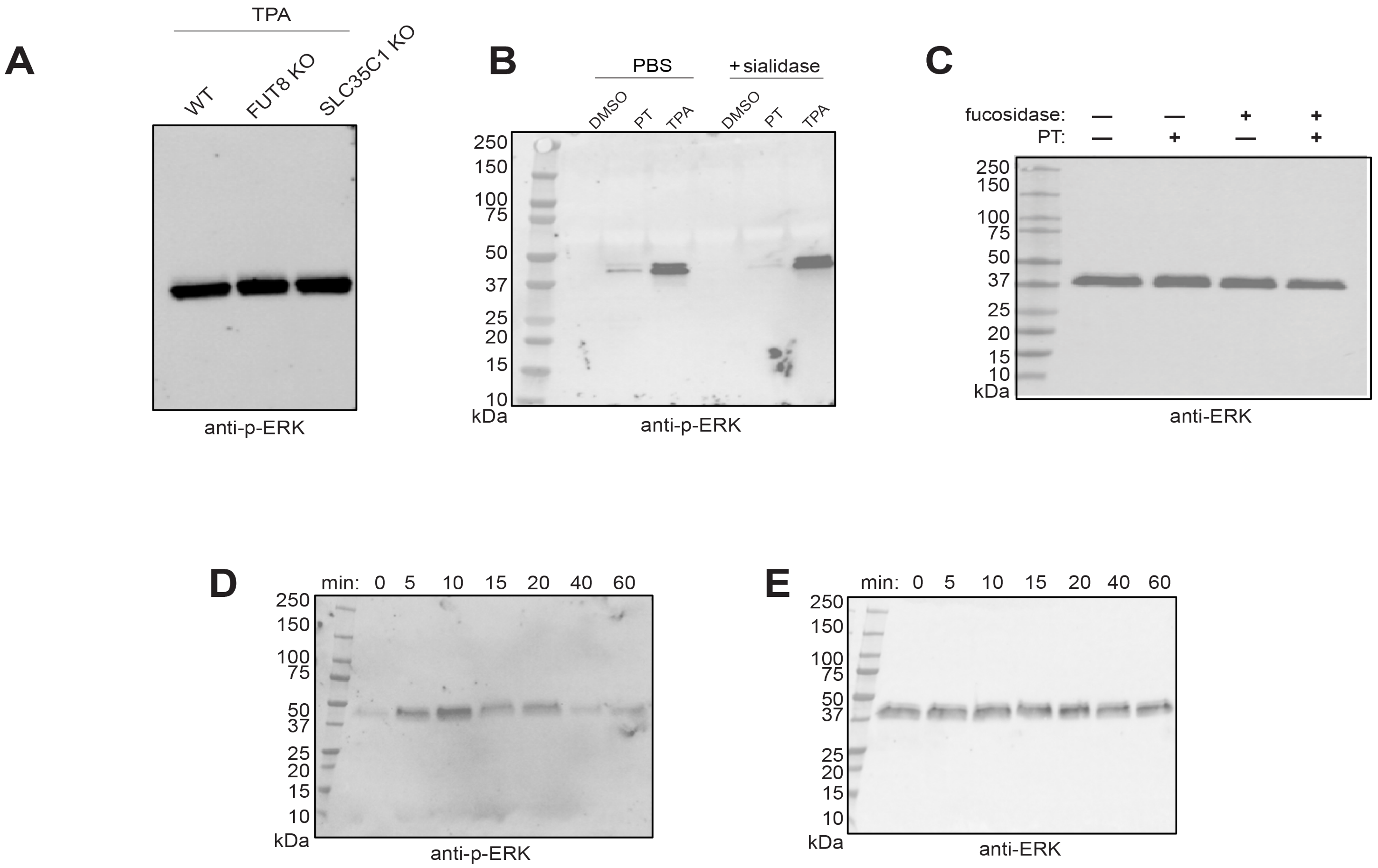


**Supplementary Figure 7. ERK phosphorylation in Jurkat cells. (A)** Phospho-ERK immunoblot of lysates from TPA-treated (60 min) WT, FUT8 KO and SLC35C1 KO Jurkat cells. **(B)** Phospho-ERK immunoblot of lysates from Jurkat cells with or without sialic acid and induced with either PT or TPA for 60 minutes. **(C)** ERK immunoblot of lysates from PT-induced (60 min) Jurkat cells treated with FucosExo. **(D)** Phospho-ERK immunoblot of lysates from PT-treated Jurkat cells at different time points, ranging from 0 to 60 minutes. **(E)** ERK immunoblot of lysates from PT-treated Jurkat cells at different time points, ranging from 0-60 minutes.
